# Selection may oppose invasion, yet favour fixation: consequences for evolutionary stability

**DOI:** 10.1101/706879

**Authors:** Chai Molina, David J. D. Earn

## Abstract

Models of evolution by natural selection often make the simplifying assumption that populations are infinitely large. In this infinite population limit, rare mutations that are selected against always go extinct, whereas in finite populations they can persist and even reach fixation. Nevertheless, for mutations of *small phenotypic effect*, it is widely believed that in *sufficiently large populations*, if selection opposes the invasion of rare mutants, then it also opposes their fixation. Here, we identify circumstances under which infinite-population models do or do not accurately predict evolutionary outcomes in large, finite populations. We show that there is no population size above which considering only invasion generally suffices: for any finite population size, there are situations in which selection opposes the invasion of mutations of arbitrarily small effect, but favours their fixation. This is not an unlikely limiting case; it can occur when fitness is a smooth function of the evolving trait, and when the selection process is biologically sensible. Nevertheless, there *are* circumstances under which opposition of invasion does imply opposition of fixation: in fact, for the *n*-player snowdrift game (a common model of cooperation) we identify sufficient conditions under which selection against rare mutants of small effect precludes their fixation—in sufficiently large populations—*for any selection process*. We also find conditions under which—no matter how large the population—the trait that fixes *depends on the selection process*, which is important because any particular selection process is only an approximation of reality.

## 1 Introduction

Adaptive dynamics is a widely used and extremely successful framework for investigating the evolution of continuous traits by natural selection (1). In this framework, it is assumed that the population is infinite and well-mixed, and that any single mutation has an extremely small phenotypic effect. One of its significant contributions is a simple method for identifying ***locally evolutionarily stable strategies*** (local ESSs). If residents are playing a local ESS then rare mutants playing a distinct but sufficiently similar strategy cannot invade the population (1, 2).

The simplicity of the notion of local ESS depends on the population being infinite. In a finite population of size *N*, selection can oppose the invasion of a mutant strategy, yet favour its fixation (*i.e*., the offspring of a single mutant will replace the resident population with probability greater than 1/*N*, the fixation probability for a neutral mutation arising in a single individual). Thus, Ref. (3) proposed a refinement of the classical definition of evolutionary stability, requiring in addition to selection opposing invasion, that selection also oppose fixation of the mutant strategy; such a strategy is said to be an ***ESS_N_*** to emphasize the finite population size.

In contrast to the conditions for evolutionary stability in finite populations, the adaptive dynamics condition for evolutionary stability does not explicitly address the possible fixation of mutants. The reason is that in an infinite, well-mixed population, a strategy that cannot invade will not fix: the effect of finitely many mutants on the residents’ fitness is “infinitely diluted” and therefore negligible (mutants can affect the residents’ fitness only if they constitute a non-negligible proportion of the population, in which case there must be infinitely many mutants). Consequently, mutants that are selected against when rare die out before they can affect residents (4). Thus, by excluding the possibility of fixation of mutants that are selected against when rare, the adaptive dynamics assumption of an infinite population inherently limits its ability to account for the effects of frequency-dependent selection.

A natural first question to ask is whether or not this limitation disappears for mutations of arbitrarily small effect: for a specific population size and selection process, if selection opposes the invasion of mutants playing a strategy sufficiently similar to the residents’, does it also opposes their fixation? In §§2 and 4, we show that the answer to this question is “*no*”, in the sense that for any population size *N*, it is possible to construct well behaved payoff functions (and a selection process) such that there is a singular strategy at which selection opposes the invasion but favours the fixation of mutations of arbitrarily small effect.

Second, given any specific game through which fitness payoffs are determined, if the population is sufficiently large and selection opposes the invasion of mutants that are sufficiently similar to residents, does it also opposes their fixation? § 5 addresses this question for the *n*-player snowdrift game—a canonical model of cooperation—and shows that this is not true in general. However, we identify a condition under which for such games, if selection opposes the invasion of sufficiently similar mutants, it generically also opposes their fixation in sufficiently large populations, regardless of the selection process.

Importantly, there is no simple rule of thumb determining how large is “large enough”; this depends on the specifics of the selection process and the game that determines fitness. That is, for any finite population, no matter how large, analysis based on adaptive dynamics—or any other framework that ignores population size and the selection process—is not sufficient in general to determine evolutionary outcomes.

## 2 Model

Our analyses are presented in the context of a standard model for the evolution of cooperation, the *n*-player snowdrift game (5–15).

### Payoffs

Members of a group of *n* individuals make costly contributions to a public good, generating a benefit (available to all group members) that depends on the total contribution made by all group members. Denoting the cost of the focal individual’s contribution *x* by *C*(*x*) and the benefit to a focal individual in a group in which the total contribution is *τ* by *B*(*τ*), the focal individual’s payoff is

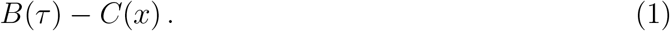

### Group formation

Groups that play the snowdrift game are formed by sampling *n* individuals uniformly and at random from the population without replacement. The population is thus well-mixed in the sense that individuals are equally likely to be sampled to play.

### Fitness

Strategies are assumed to be inherited and therefore subject to selection. For simplicity, we assume that individuals play many rounds of the game between reproductive events (*i.e*., at each time step) and that their fitnesses are simply their average payoffs (1) from these games.

### Group and population sizes

We assume that the group size *n* ≥ 2. (If there were only one individual in a “group” then its optimal strategy would not depend on the behaviour of others and evolutionary game-theoretic considerations would be irrelevant.) The population can be infinite (*N* = ∞); if it is finite, we assume that it is larger than the group size, *i.e., N* > *n*; in particular, *N* ≥ 3. (If the entire population were to play the game together, *i.e*., for *n* = *N*, individuals contributing the least would always have the highest fitness, so populations would inevitably evolve to defection, *i.e*., contributing nothing.)

### Trait Substitution

Mutations are assumed to be sufficiently rare that no more than two strategies are present in the population at any time.

## 3 Strategy dynamics

### 3.1 Infinite populations

When the snowdrift game described above is played in an infinite population (*N* = ∞), the evolution of strategies (*i.e*., contributions to the public good) is well-described by the canonical equation of adaptive dynamics (16). A strategy *X* is ***evolutionarily stable*** if (1, 2)

1. it is ***singular***, *i.e*., directional selection vanishes in its vicinity (or more precisely, the local fitness gradient vanishes as it is approached),

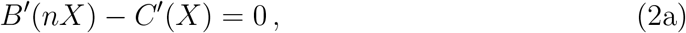

and
2. selection ***opposes invasion*** of mutants playing an arbitrarily similar strategy (which is ensured by requiring that the fitness of an invading mutant as a function of the mutant strategy is concave when mutants play the resident strategy)

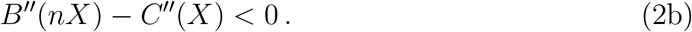

### 3.2 Finite populations

To find evolutionary outcomes of the *n*-player snowdrift game (§ 2) when played in a finite population (*N* < ∞), we use a framework for analyzing evolutionary stability in finite populations introduced in Ref. (17). We denote by 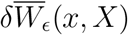 the ***difference in mean fitness*** of mutants (playing *x*) and residents (playing *X*), where *ϵ* is the proportion of the population playing the mutant strategy *x* (so *ϵ* takes one of the values 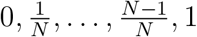).

Similar to the infinite-population case, selection opposes invasion of a population of residents playing *X* by mutants playing (an arbitrarily similar strategy) *x* if the expected fitness of such mutants is lower than for residents, that is, 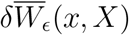 attains a local maximum as a function of *x* for *x* = *X*. This occurs if the following two conditions hold.

1. The resident strategy *X* is ***singular***, *i.e*., directional selection vanishes in its vicinity, or more precisely, the local fitness difference gradient vanishes as it is approached, *i.e*., 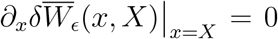; see definition 4.3.5 in Ref. (17). For the snowdrift game, the condition for a singular strategy can be written

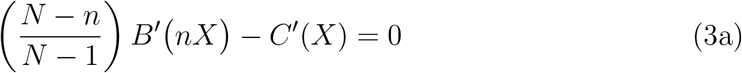

[*cf*. equation (4.64) in Ref. (17)].
2. The fitness of an invading mutant as a function of the mutant strategy is concave when mutants play the resident strategy, 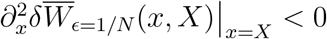. For the snowdrift game, this concavity condition can be written

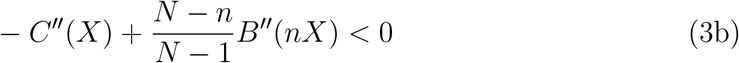

[*cf*. equations (4.64) and (4.71) in Ref. (17)].

Equation (3a) is a necessary condition for selection opposing invasion of mutants playing strategies sufficiently similar to the residents: if (3a) does *not* hold, then the fitness difference between the invading mutant and residents is either increasing or decreasing as a function of the mutant’s strategy; in the increasing (resp. decreasing) case, invading mutants contributing slightly more (less) than the residents obtain higher fitness than the residents^*^.

In a finite population (*N* < ∞), the assumptions that define the our model framework (§ 2) do not completely determine the strategy dynamics that unfold following the introduction of a mutant. In particular, fixation probabilities naturally depend on the ***selection process***, *i.e*., on how fitnesses are used to determine changes in the frequencies of the two traits that are present in the population over time [see Ref. (18)]. Without specifying a particular selection process, it is in general impossible to identify strategies that are ESS_N_s (*i.e*., evolutionarily stable in a population of size *N*); whether selection opposes the *fixation* of mutants playing a strategy sufficiently similar to the residents depends on the selection process.

In the next two subsections, we introduce further notation related to the mean fitness difference, and a class of selection processes that we will use in later sections.

#### 3.2.1 Curvatures of the mean fitness difference

We now introduce convenient notation to simplify the Taylor expansion of the mean fitness difference 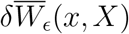 in the mutant strategy *x* about a singular resident strategy *X*.

Using equation (3a) and the identity 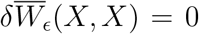 (neutral mutations do not confer a fitness advantage), for any number of mutants *i* (1 ≤ *i* ≤ *N* − 1) we can write the mean fitness difference

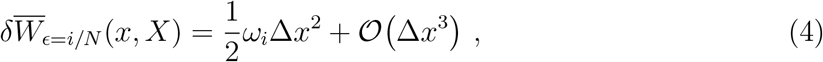

where Δ*x* = *x* − *X*, and

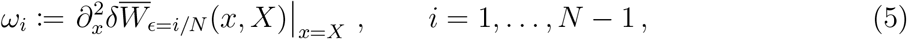

are the ***fitness difference curvatures***. Note that *ω_i_* depends on the resident strategy *X*, but we make this dependence implicit for notational convenience.

Using equations (4.64) and (4.71) of Ref. (17), the coefficient *ω_i_* is given by

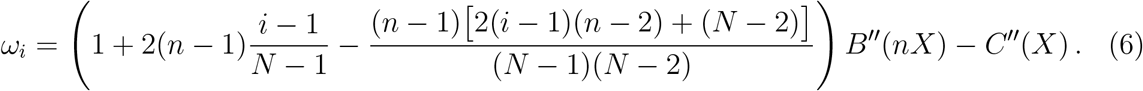

In particular,

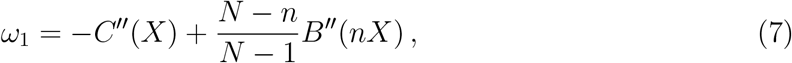

so setting

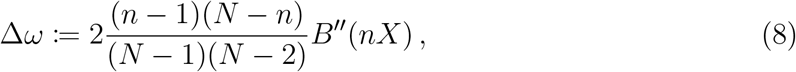

we have

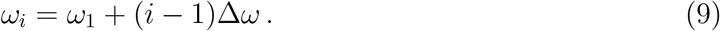

#### 3.2.2 Symmetric birth-death processes

Some of our analysis involves identifying situations in which selection favours fixation. To that end, in appendix A we define a class of biologically sensible selection processes—which we call ***symmetric birth-death (or SBD) processes***—for which fixation probabilities can be conveniently expressed in terms of differences in mean fitness. If there are *i* mutants in the population (with 1 ≤ *i* ≤ *N* − 1), and if we denote the mean fitness difference by

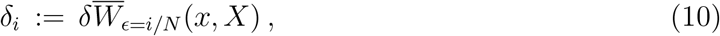

then the inverse of the fixation probability is

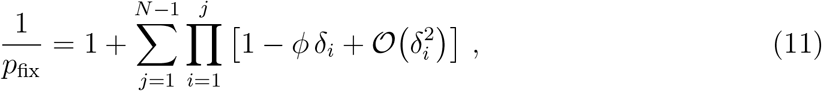

where *ϕ* > 0 is a parameter that depends on which SBD process is chosen. SBD processes are used in § 4 for analyses that depend only on equation (11) and in § 6.2 for numerical simulations; the particulars of how SBD processes are defined (in appendix A) are not essential to understand the results. In § 5 and § 6.3, we present more general results that are not specific to SBD processes.

## 4 Selection can oppose invasion but favour fixation of arbitrarily similar mutants

In this section, we demonstrate that for any given population size *N* and any group size *n* < *N*, there are games for which selection opposes invasion but favours fixation (of mutant strategies that can be arbitrarily close to a singular strategy played by residents).

Consider the evolution of contributions to an *n*-player snowdrift game (as described in § 2) in a finite population of size *N* governed by an SBD selection process (defined in appendix A). In this situation, we show that it is possible to find benefit and cost functions, *B* and *C*, and a resident strategy *X*, such that

> mutants that play a strategy (*x*) that is different from—but sufficiently similar to—the resident strategy (*X*) obtain *lower fitness when rare*, yet *selection favours the fixation of such mutants*.

The conditions for this are stated precisely in proposition 1, which we prove in appendix B.

### Proposition 1.

*Consider an evolving population of finite size N, where fitnesses are determined by playing the n-player snowdrift game as described in § 2. If residents play a singular strategy* (i.e., *a strategy X that satisfies* *equation* (3a)), *and in addition*,

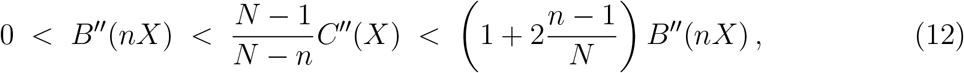

*then for any sufficiently similar strategies x* (i.e., *for* |*x* − *X*| *sufficiently small*), *selection opposes the invasion of mutants playing x, but favours their fixation under any SBD selection process (§ 3.2.2 and appendix A)*.

It is easy to find functions *B* and *C* that satisfy the conditions in proposition 1. In § 6.2, we construct explicit examples of games that satisfy the hypotheses of proposition 1 and therefore exhibit fixation of strategies that are opposed by selection when rare.

## 5 Evolutionary outcomes can depend on the selection process, even in large populations

In § 4 we fixed the *population size N* and found conditions guaranteeing that selection opposes invasion but favours fixation; these conditions are satisfied by many snowdrift games. In this section, we fix the *game* (a snowdrift game with specific benefit and cost functions and a fixed group size^†^) but *not* the population size. We consider situations in which the game has an ESS if played in an infinite population and ask whether it also has an ESS_N_ if played in a sufficiently large finite population.

Proposition 2 below (proved in appendix C) provides the answer, which is not as simple to state as one might hope. The existence of an infinite-population singular strategy 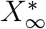 generically implies the existence of a finite-population singular strategy 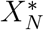 when the same game is played in a finite population of sufficiently large size *N*; in fact, 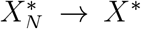 as *N* → ∞. We identify a condition (namely, condition (16)) guaranteeing that for sufficiently large population size *N*, 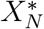 is a (local) ***universal ESS***_*N*_ (UESS_N_), that is, selection opposes the invasion and fixation of mutations of arbitrarily small effect, *regardless of the selection process*. However, if condition (16) does not hold, then the evolutionary stability of 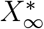 does not generally imply that 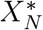 is evolutionarily stable: we identify a condition under which, when residents play 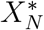, selection may either favour or oppose the fixation of mutations of arbitrarily small effect, *depending on the selection process*. In §6.3, we also construct explicit examples of snowdrift games that exhibit this behaviour.

### Proposition 2.

*In a snowdrift game as defined in § 2, suppose the benefit and cost functions, B and C, and the group size n, are such that there is a strategy* 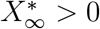 *satisfying the adaptive dynamics conditions for evolutionary stability* (2), *and at which*

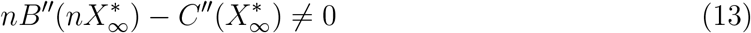

(*which holds generically*^‡^). *If this same game is played in a finite population of sufficiently large size N, then for each such N there is a singular strategy* 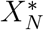, *and*

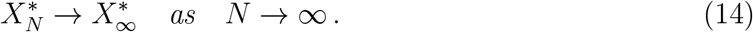

*If, in addition, the fitness difference curvature ω*_*N*−1_ (*equation* (5)) *is negative for sufficiently large N*, i.e., *if there exists a population size <u>N</u> such that*

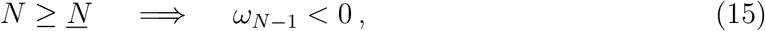

*then there exists <u>N</u>*\* ≥ *<u>N</u> such that for any N* ≥ *<u>N</u>*\*, 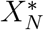 *is a UESS_N_. A sufficient condition for such an <u>N</u> to exist is that*

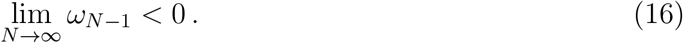

*Conversely, if there exists* 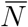 *such that*

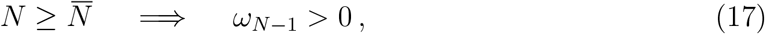

*then there exists* 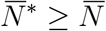 *such that for all* 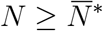, *for mutations of arbitrarily small effect, selection favours fixation for some selection processes, but opposes fixation for other selection processes; a sufficient condition for such an* 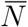 *to exist is that*

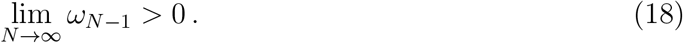

Conditions (16) and (18) are easy to check because the limit can be expressed directly in terms of the benefit and cost functions: Equations (8) and (9) give

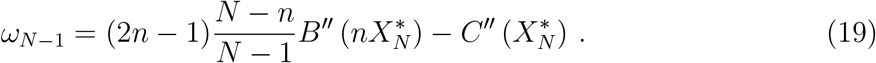

Since, in addition, 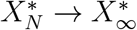 as *N* → ∞ (14), we have

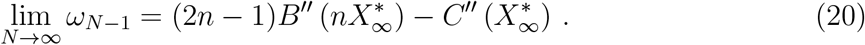

Note that condition (15) [condition (17)] is more general than condition (16) [condition (18)]: the sign of lim_*N*→∞_ *ω*_*N*−1_ being negative (positive) is not necessary for the existence of 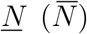, because it is possible that *ω*_*N*−1_ < 0 (> 0) for all sufficiently large *N*, but that lim_*N*→∞_ *ω*_*N*−1_ = 0. However, equation (20) implies that lim_*N*→∞_ *ω*_*N*−1_ exists and vanishes 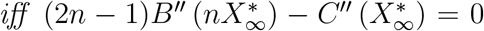, which generically is *not* satisfied. In other words, generically, either condition (16) or condition (18) holds.

In the unlikely situation that *neither* condition (15) *nor* (17) holds (which can only happen if lim_*N*→∞_ *ω*_*N*−1_ = 0), there are two possibilities:

- There are increasing, unbounded sequences 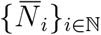 and {*<u>N</u>_i_*}_*i*∈ℕ_ such that *ω*_*<u>N</u>_i_*−1_ < 0 and 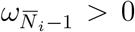 for all *i* ∈ ℕ: in this case, corollary 5.4 and lemma 4.6 in Ref. (18) (respectively) imply that 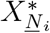. is a UESS_N_ and 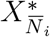 is *not* a UESS_N_ for all *i* ∈ ℕ. For any 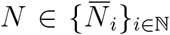, if residents play 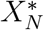, some selection processes favour fixation of mutations of arbitrarily small effect, while other selection processes oppose their fixation.
- The fitness difference curvature *ω*_*N*−1_ vanishes for all sufficiently large *N* (*i.e*., there exists *N*_0_ such that *ω*_*N*−1_ = 0 for all *N* > *N*_0_): in this case, it is possible that 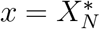 is a minimum, maximum or inflection point of 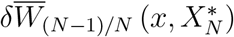 for all sufficiently large *N*. Consequently, if *ω*_1_ < 0 it is still possible that 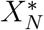 is a UESS_N_; but, it is also possible that some selection processes favour fixation of mutations of arbitrarily small effect, while other selection processes oppose their fixation.

## 6 Examples

In this section, we illustrate the predictions of propositions 1 and 2 with examples, using a subclass of snowdrift games that we define in § 6.1. The particular examples are then described in §§6.2 and 6.3.

### 6.1 A class of quadratic snowdrift games

Consider a snowdrift game (§ 2) with quadratic benefit and cost functions (similar to (5)),

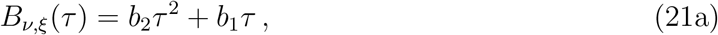

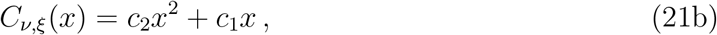

where the coefficients are

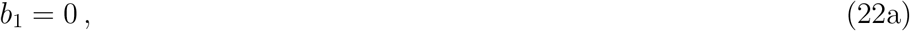

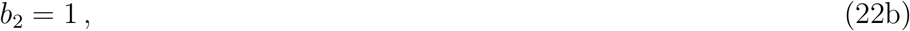

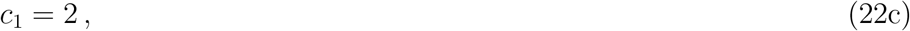

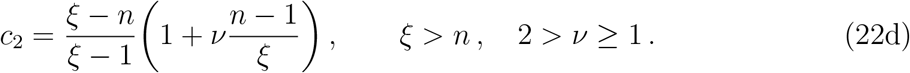

We denote such a game for particular *ν* and *ξ* as 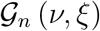, and the family of all such games for fixed group size *n* as

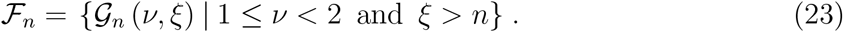

Note that games in this class differ only in their cost functions.

#### 6.1.1 Singular strategies

When snowdrift games with quadratic benefit and cost functions [equation (21)] are played in an infinite population, solving equation (2a) gives the unique singular strategy,

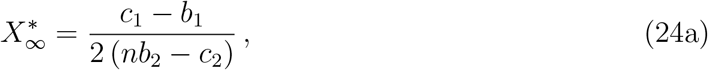

whereas when they are played in a finite population of size *N* > *n*, equation (3a) gives the singular strategy,

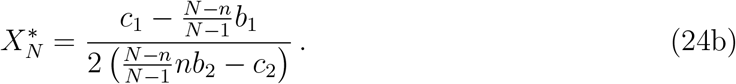

Note, however, that for some choices of the coefficients (*b*_1_, *b*_2_, *c*_1_, *c*_2_), 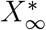 and 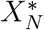 can be negative (and therefore biologically irrelevant). For the specific benefit and cost coefficients given by equation (22), *i.e*., for all games in the class 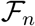, equations (24a) and (24b) become

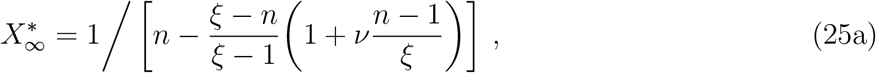

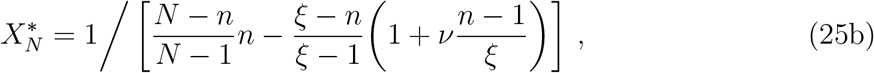

It is straightforward to show that 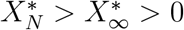 for any *ξ* > *n* and *ν* ∈ [1, 2).

#### 6.1.2 Sufficient condition for evolutionary stability in an infinite population

To guarantee that the singular strategy 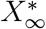 given by equation (25a) is evolutionarily stable when played in an infinite population, a sufficient condition is that

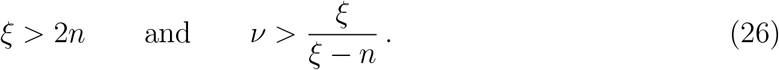

To see this, we verify that condition (26) implies that condition (2b) is satisfied: With quadratic benefit and cost functions, condition (2b) yields *c*_2_ > *b*_2_, and inserting equation (22) gives

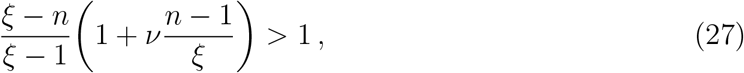

which simplifies to

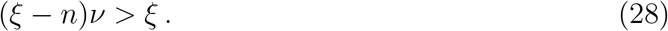

This is equivalent to condition (26) if *ξ* > 2*n*.

### 6.2 Evolutionary games with different outcomes in finite populations and infinite populations

Given a finite population size *N*, we now consider the subclass of games 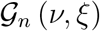 for which *ξ* = *N, i.e*., 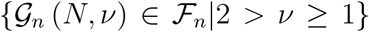. Although 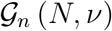 is parameterized using the given *N*, note that the games in this subclass can also be played in an infinite population.

When a game in 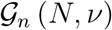 is played in a finite population of size *N*, proposition 1 applies. Thus, under an SBD selection process (appendix A), if residents play the singular strategy 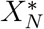, selection favours fixation of mutations of arbitrarily small effect, so 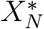 is not an ESS_N_. Since cooperative strategies (*X* > 0) that are not singular cannot be ESS_N_s, a game 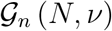 does not have a cooperative ESS_N_ when played in a population of size *N* under any SBD process. By contrast, if the same game 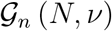 is played in an infinite population, then 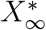 [given below in equation (29a)] is an ESS for any *ξ* = *N* > 2*n* (see § 6.1.2). We corroborate this prediction using individual-based simulations in figure 1.

**Figure 1:**
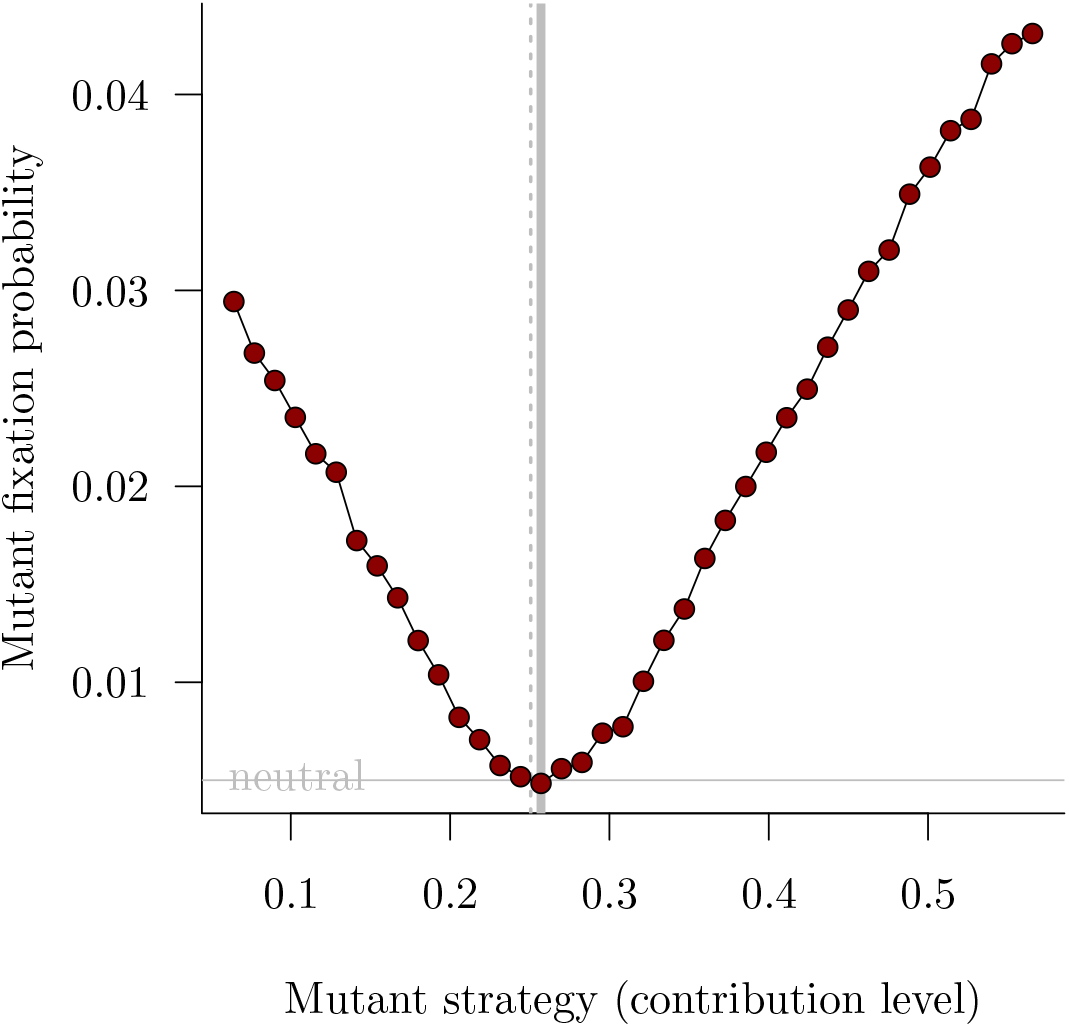
Selection opposing invasion but favouring fixation in a quadratic snowdrift game (§ 6.1; *ν* = 3/2, *ξ* = 200) with group size *n* = 5 and total population size *N* = 200. The finite-population singular strategy 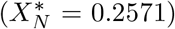 is shown with a thick vertical grey line. The associated infinitepopulation ESS 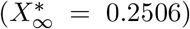 is shown with a thin dotted vertical grey line. The fixation probability of a neutral mutation (1/*N* = 0.005) is shown with a horizontal grey line. The red dots show the fixation probability of mutants when residents play the ESS_N_, based on 10^5^ simulations for each mutant strategy, under a symmetric birth-death (SBD) selection process with transition probability ratio *R*(*x*) = *e*^−*x*^ (so *ϕ* = 1 in equation (11); see equation (41) in appendix A).

To verify that proposition 1 holds for any game 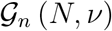 (with 2 > *ν* ≥ 1), note first that when *ξ* = *N*, equation (25a) reduces to

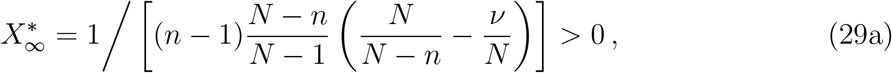

and equation (25b) becomes

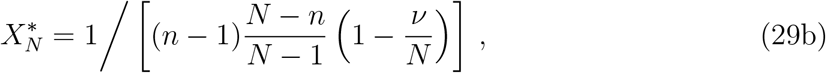

which is positive for any *N* ≥ 2 because 2 > *ν* > 0.

Next, substituting equation (21) into condition (12) gives

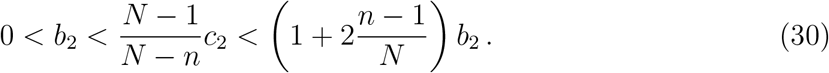

For the specific coefficients of the benefit and cost functions given by equation (22), condition (30) becomes

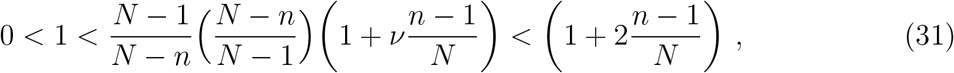

which manifestly holds for *ν* ∈ [1, 2).

### 6.3 Games for which evolutionary outcomes differ between infinite and arbitrarily large but finite populations

We now apply proposition 2 to identify games in the class 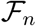 that have an ESS when played in an infinite population but—depending on the selection process—either have, or do not have, an ESS_N_ when played in arbitrarily large finite populations^§^. To do this, we must find games 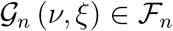 that (i) have an infinite population ESS 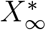, (ii) satisfy condition (13), and (iii) have the property that there is a population size 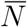 such that condition (17) is satisfied.

First, to ensure that there is always an infinite population ESS, we assume *ν* and *ξ* satisfy condition (26).

Second, we note that for games in 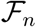, condition (13) simplifies to

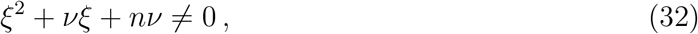

which holds because *v, ξ* and *n* are all positive. Hence condition (32) [and therefore condition (13)] holds for any *ν* ∈ [1, 2).

Finally, we find that with 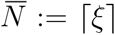, condition (17) is satisfied: Inserting equation (22) in equation (19), we have

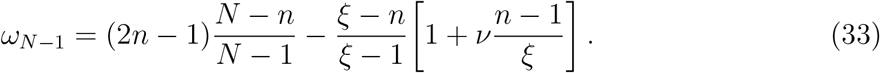

Substituting *ξ* = *N* in equation (33) gives

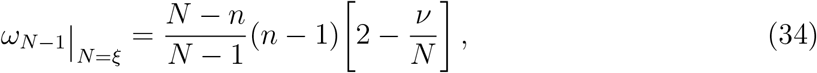

which is positive because *N* > 1 and 2 > *ν* > 0. Since (*N* − *n*)/(*N* − 1) increases with *N*, so does *ω*_*N*−1_, and hence

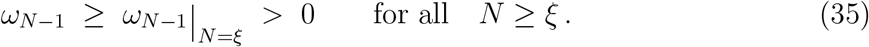

## 7 Conclusion

Evolutionary game theory has been developed primarily under the approximation of an infinite background population (2, 5, 19–23). In this setting, the notion of evolutionary stability can be formalized simply as “selection opposes invasion” and the details of the selection process are irrelevant. In finite populations, evolutionary stability requires the additional condition that “selection opposes fixation”, and which strategies are stable (ESS_N_s) depends, in general, on the selection process (18).

The traditional justification for the infinite population approximation is that sufficiently large finite populations behave as if they were infinite (4, §2.1). Here, we have challenged this conventional wisdom by demonstrating two mechanisms by which inferences drawn from evolutionary games played in an infinite population can turn out to be incorrect for more realistic, finite populations. First, we have shown that for any finite population size, there are biologically sensible situations in which selection favours the fixation of mutants, even though selection opposes their invasion when rare (proposition 1; example in § 6.2). Second, we have identified conditions on *n*-player snowdrift games such that an infinite-population cooperative ESS exists, yet in a finite population—no matter how large—the existence of a cooperative ESS_N_ depends on the selection process, *i.e*., a cooperative ESS_N_ exists under some selection processes but not others (proposition 2; example in § 6.3).

Thus, while the infinite population approximation is convenient mathematically and leads to simpler predictions, those predictions can be misleading in finite populations (no matter how large). Infinite-population models should, therefore, be applied carefully and cautiously. To that end, we have also identified conditions under which the infinite population approximation *does correctly predict* evolutionary outcomes in sufficiently large finite populations. More precisely, Proposition 2 formalizes conditions—on snowdrift games—that guarantee that if an infinite-population ESS exists then in sufficiently large (but finite) populations there is a strategy that is *universally* evolutionarily stable (*i.e*., is an ESS_N_ for any selection process). Under these conditions, the adaptive dynamics framework is useful and correctly predicts evolutionary outcomes in sufficiently large (finite) populations.

## Acknowledgments

DE was supported by the Natural Sciences and Engineering Research Council of Canada (NSERC). CM was supported by the United States Defense Advanced Research Project Agency NGS2 program (grant no. D17AC00005), and the Army Research Office (grant no. W911NF1810325).

## APPENDICES

## A Symmetric birth-death selection processes

Let *M*_p_(*t*) be the number of mutant individuals at time *t*; since individuals are either mutants or residents, *M*_p_(*t*) completely specifies the ***population state*** at time *t*. If *M*_p_(*t*) = *i* for 1 ≤ *i* ≤ *N* − 1 then both mutants and residents are present so we refer to a ***mixed population state***.

We define a discrete-time birth-death process that—based on the fitness difference between mutants and residents—changes the composition of the population over time; for convenience, we use *P* to denote both this process, and the transition matrix that defines it. Specifically, we set

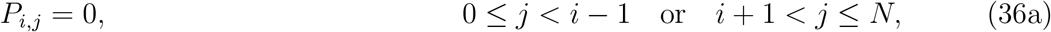

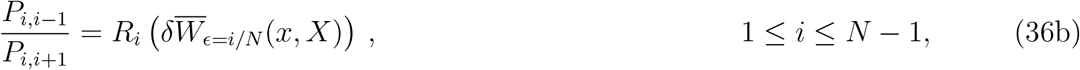

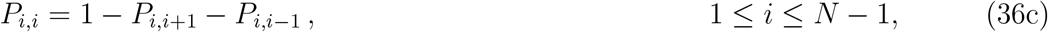

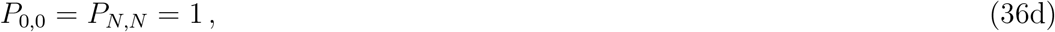

where the dependence of the ***transition probability ratios*** *R_i_* (*i* = 1,…, *N* − 1) on the mean fitness differences 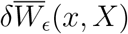 (§ 3.2) will be specified shortly. The following two conditions must be satisfied for equation (36b) to make biological and mathematical sense:

a. The right hand side of equation (36b) must be non-negative for any fitness difference 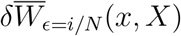 (otherwise some transition probabilities will be negative). Moreover, if 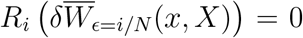 for some *i* then mutants cannot go extinct (fixation of the mutants is certain) so we assume *R_i_*(·) > 0. [The consistency condition (b) stated next also independently excludes the possibility that 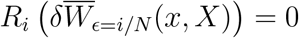.]
b. The transition probability ratios *R_i_* must satisfy a consistency condition. If a population consists of *i* and *N* − *i* individuals of types *A* and *B*, respectively, then the ratio of the probabilities that the number of individuals of type *A* increases, and decreases, must be independent of whether type *A* is labelled as the mutant or resident. Mathematically, if mutants and residents are interchanged, state *i* becomes state *N* − *i*, and the mean fitness difference 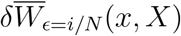 becomes 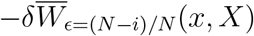. Thus, we require that

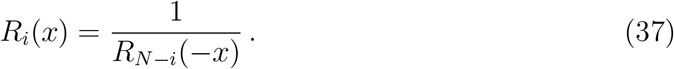
c. The transition probability ratios *R_i_* must be decreasing functions of the fitness difference. This assumption is motivated by the biologically sensible intuition that if the mutants have a fitness advantage over the residents, increasing this advantage should increase the probability that the number of mutants increases (and lower the probability that the number of mutants decreases). Conversely, if the residents have a fitness advantage over the mutants (*i.e*., 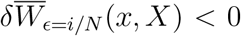) then increasing this advantage should *decrease* the probability that the number of mutants increases.

For simplicity, we assume that the ratios of probabilities of mutants increasing and decreasing (*P*_*i,i*−1_/*P*_*i,i*+1_) depend on the mean fitness difference 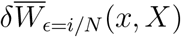, but not on the population state *i*, so that, with minor abuse of notation, only one transition probability ratio function is needed, *R_i_* = *R* for all *i* = 1,…, *N* − 1, and

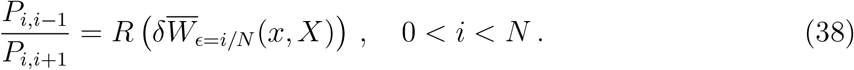

Condition (37) then becomes *R*(*x*)*R*(−*x*) = 1. Since *R* > 0, this is equivalent to

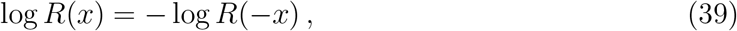

so *r*(*x*) := log *R*(*x*) is an odd function. It follows that the transition probability ratio must be of the form

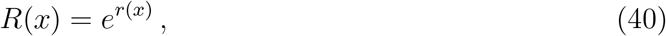

where *r* : ℝ → ℝ is odd. Since *R* is decreasing, *r* must also be decreasing.

We assume henceforth that *r* is analytic in a neighbourhood of *x* = 0. Because *r* is odd, *r*^(*n*)^(0) = 0 for any non-negative even integer *n*, and letting *ϕ* = −*r′*(0) > 0, we have

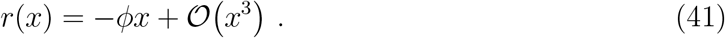

Consequently, for *i* = 1,…, *N* − 1, equation (36b) gives

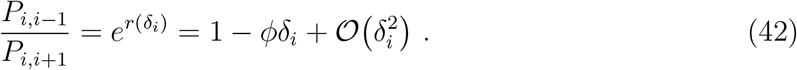

where 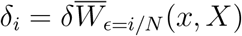 as in equation (10).

Equation (36) leaves some freedom in that we do not specify how likely it is for the population to remain at the same state (*i.e*., that *M*_p_(*t* + 1) = *M*_p_(*t*)); this affects the speed of evolution, but not the fixation probabilities. For concreteness, we can set *P_i,i_* = 0 for all *i* = 1,…, *N*, in which case equations (36b) and (36c) can be solved explicitly to obtain

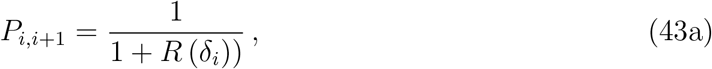

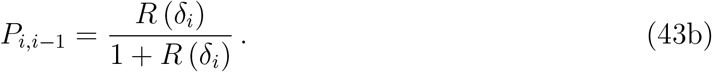

We have thus constructed a class of birth-death processes, determined by the choice of the logarithm *r* of the transition probability ratio *R*, which must be a decreasing odd function (taking *r*(*x*) = −*x* yields the simplest such process). These processes *P* are well-behaved in the sense that for any resident and mutant strategies

I. at any time *t* the number of individuals of the type with higher fitness is expected to increase in the next time step,
II. no new mutations are introduced, so that monomorphic populations are absorbing states of *P*, and
III. starting from any mixed population state it is possible for the mutation to either fixate or become extinct, that is, the probabilities of these outcomes happening at some future time are nonzero.

Consequently, *P* is indeed a selection process as defined in Ref. (18). Moreover, *P* satisfies the consistency condition (37) by construction and, for smooth *r*, depends smoothly on the mean fitness difference 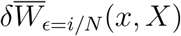.

If initially one mutant individual playing *x* enters the population (*M*_p_(0) = 1), the probability that the mutation fixes can be calculated exactly (because *P* is a birth-death process) and is given by

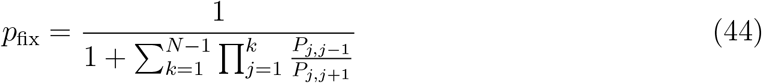

[see, for example, Appendix C of Ref. (18) for a detailed proof]. Inserting equation (42), we obtain equation (11).

## B Proof of proposition 1

Let *X* be a singular strategy (equation (3a)) for which condition (3b) holds, so that selection opposes invasion of a population of residents playing *X* by mutants playing *x* sufficiently close to *X*. We aim to find additional conditions on the benefit and cost functions (*B* and *C*) such that, for SBD processes (§ 3.2.2), selection *favours* the fixation of *x* sufficiently near *X*, even though it opposes invasion of *x* when rare. Denoting the fixation probability of the mutant strategy when one mutant individual initially enters the population by *p*_fix_ we wish to find conditions under which *p*_fix_ > 1/*N*.

First, observe that from equation (4), if *ω_i_* < 0, then 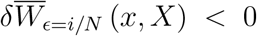 for all *x* sufficiently close to but different from *X*, and conversely, if *ω_i_* > 0, then 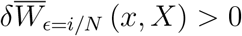 for such mutants. Thus, if *ω_i_* < 0 for all *i* = 1,…, *N* − 1, then 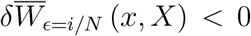 for all *x* sufficiently close to but different from *X*, so corollary 5.4 in Ref. (18) implies that selection opposes fixation of such mutants regardless of the selection process. Thus, in order for selection to favour the fixation of mutants playing *x* close to *X*, there must be some number of mutants *i* (1 ≤ *i* ≤ *N* − 1) for which *ω_i_* > 0.

The definition of the fitness difference curvatures *ω_i_* [equation (5)] implies that condition (3b) is equivalent to *ω*_1_ < 0. Therefore, to stack the odds in favour of the mutants fixing, we will require

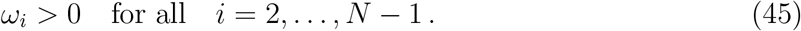

Because *ω_i_* is linear in *i* [equation (9)], this is achieved if

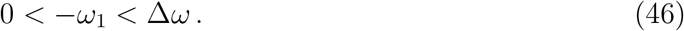

Using equations (7) and (8), this is equivalent to

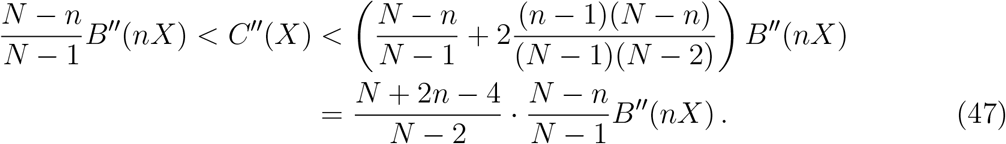

Henceforth, we assume that condition (46) [or equivalently, condition (47)] holds. Then, Δ*ω* > 0, so the curvatures *ω_i_* increase with *i*. Thus, we can bound fixation probabilities for invading mutants under an SBD process (§ 3.2.2) by substituting equation (4) into equation (11) to get

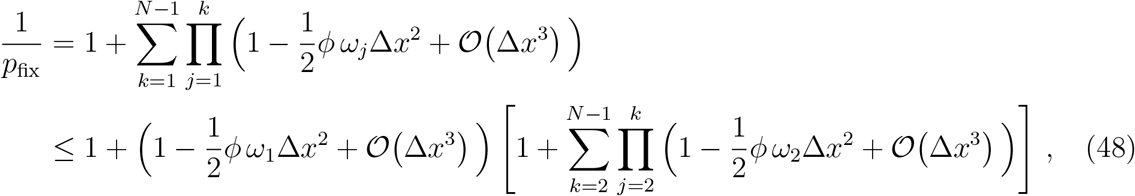

where we have used *ϕ* > 0 [equation (41)] and 0 < *ω*_2_ ≤ *ω*_3_ ≤ ⋯ ≤ *ω*_*N*−1_ to obtain inequality (48). Simplifying the term in square brackets gives

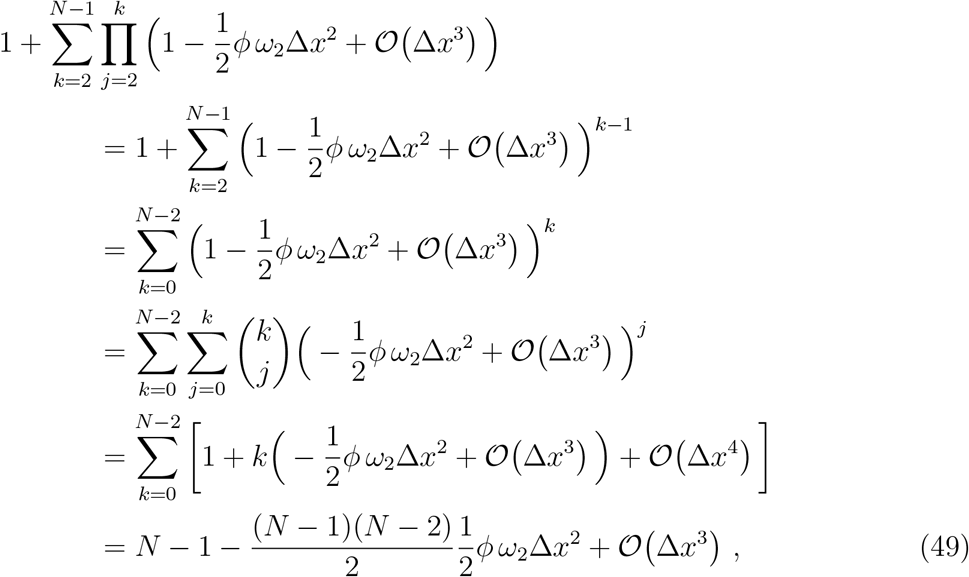

and hence

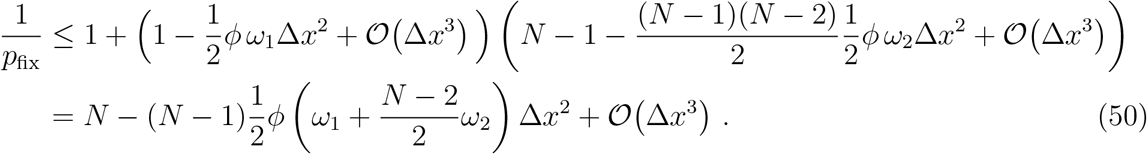

It follows that if Δ*x* is sufficiently small then 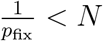 (so 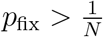 as desired) provided that the coefficient of Δ*x*^2^ in equation (50) is negative, *i.e*., provided that

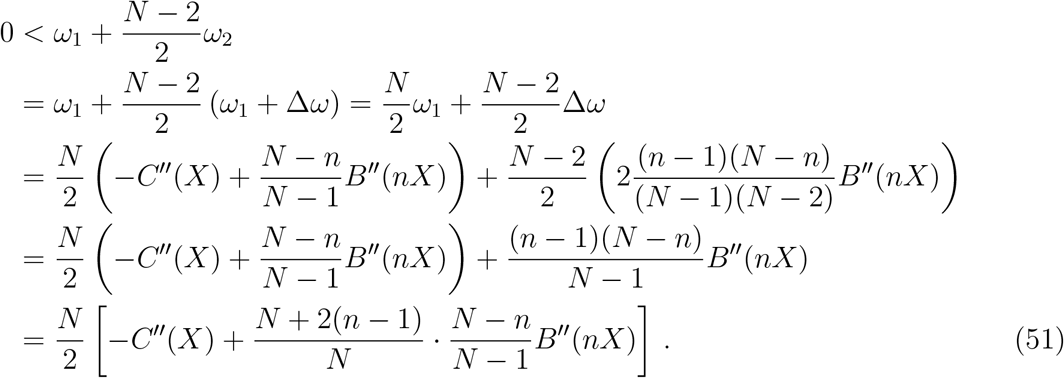

Thus, 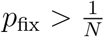 if

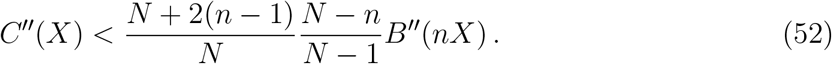

Now recall that in order to ensure that selection opposes invasion of similar mutants, but that the mean fitness of mutants is higher than that of residents when the population contains two or more mutants, condition (47) must hold. Therefore, all that remains is to determine is under what circumstances equations (47) and (52) both hold. Because *N* > *n* ≥ 2, we have

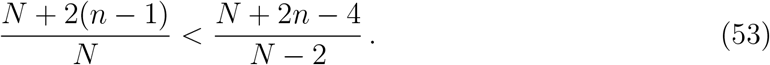

Furthermore, condition (47) also implies

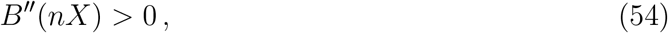

so we have

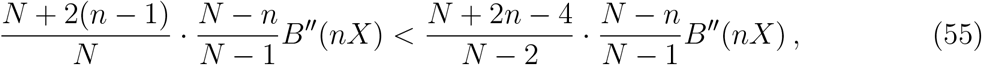

that is, the right hand side of equation (52) is smaller than that of condition (47). Hence, both equations (47) and (52) are satisfied if

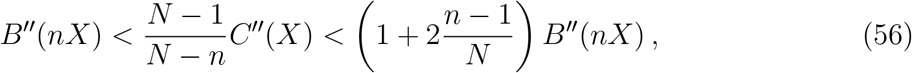

in which case both *ω*_1_ < 0 and 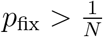, as desired^¶^.

## C Proof of proposition 2

It can be shown that if the population size *N* is sufficiently large, there is a finite-population singular strategy 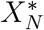 [*i.e*., a solution of equation (3a)]; moreover, the sequence of these singular strategies approaches 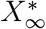 as *N* → ∞ (see lemma 3 in appendix D). Henceforth, assume without loss of generality that *N* is sufficiently large that the singular strategy 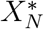 exists. Taking the limit *N* → ∞ in equation (7) we find

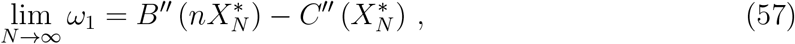

which is negative because we assume that equation (2) is satisfied. Consequently, for *N* large enough, *ω*_1_ < 0.

Now, suppose that *ω*_*N*−1_ < 0 for all *N* > *<u>N</u>*. Then, there exists *<u>N</u>*\* ≥ *<u>N</u>* such that if *N* ≥ *<u>N</u>*\* then both *ω*_1_ and *ω*_*N*−1_ are negative. Since *ω_i_* is linear in *i* [equation (9)], this implies that *ω_i_* < 0 for all *i* = 1,…, *N* − 1. Hence, from equation (4), we have 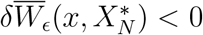 for *x* sufficiently close to but different from 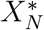. In other words, if residents play 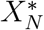, mutants playing a strategy that is sufficiently similar to the residents’ obtain a lower fitness than residents, regardless of the number of mutants (*i.e*., for all *i* = 1,…, *N* − 1). Thus, selection opposes both invasion and fixation of such mutants regardless of the selection process (18, corollary 5.4), so 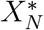 is a UESS_N_.

Similarly, if *ω*_*N*−1_ > 0 for all 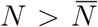, then there exists 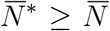 such that *ω*_1_ < 0 and *ω*_*N*−1_ > 0, so *ω_i_* changes sign as a function of *i*. Consequently, for all 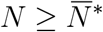, when mutants play strategies that are arbitrarily similar to the residents, the mean fitness difference 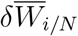 is positive for some numbers of mutants and negative for others. Consequently, from Lemma 4.6 in Ref. (18), it follows that selection favours fixation of such mutants for some selection processes, and selection opposes their fixation for other selection processes.

## D Existence and convergence of finite-population singular strategies

Lemma 3 below shows that the existence of a singular strategy when a game is played in an infinite population generically implies the existence of a corresponding singular strategy when it is played in a sufficiently large finite population.

### Lemma 3

(Existence of singular strategies in sufficiently large populations). *Consider an evolving population of finite size N in which fitness is determined by payoffs from an n-player snowdrift game (§ 2) for which the second derivatives of the benefit (B) and cost (C) functions exist. Suppose that a singular strategy* 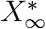 *exists when the same snowdrift game is played in an infinite population, and that* 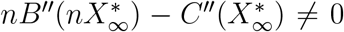 *(which holds generically). Then, if the population size N is sufficiently large, a corresponding singular strategy* 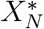 *exists. Furthermore, the sequence of finite-population singular strategies* 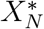 *that result from playing this game in finite populations of different (sufficiently large) size approaches* 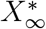 *as N* → ∞.

*Proof*. Letting

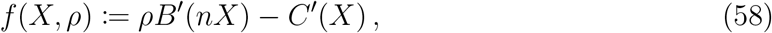

we have 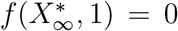, because 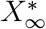 is singular in an infinite population, and so satisfies equation (2a). Noting that 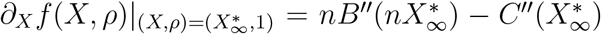, the hypothesis that 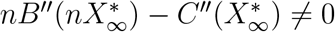 implies that

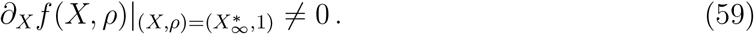

Therefore, from the implicit function theorem [*e.g*., (24, Theorem 12.40)], there is a differentiable function *X*(*ρ*) defined in a neighbourhood of *ρ* = 1, such that

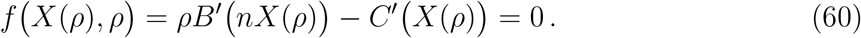

Now define

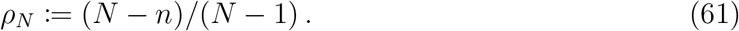

Since *ρ_N_* → 1, it follows that for sufficiently large population size *N, f*(*X*(*ρ_N_*), *ρ*) = 0 can be solved implicitly to yield 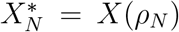. These solutions 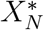 are singular [*i.e*., solve equation (3a)]. Moreover, 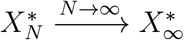 because *X*(*ρ*) is continuous.

## Footnotes

* Moreover, because 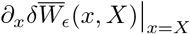 is independent of the proportion of mutants in the population, mutants that obtain a higher fitness when rare do so regardless of their frequency in the population, so by Corollary 5.4 of (18), selection favours their fixation.

† By a fixed group size we mean that the size of the interacting group (*n*) is independent of the population size (*N*). For example, the typical size of groups travelling in cars that are obstructed by snowdrifts would be the same in small and large cities.

‡ In particular, in the biologically sensible case of accelerating costs and decelerating benefits, condition (13) always holds.

§ For sufficiently large populations, condition (12) does not hold for quadratic snowdrift games, so proposition 1 does not apply to these games. Consequently, we cannot say whether or not an ESS_N_ exists when such games are played in arbitrarily large but finite populations under SBD selection processes.

¶ Because 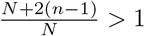, the right hand side of condition (56) is greater than the left whenever *B″*(*nX*) > 0, so condition (56) cannot be satisfied if the benefit function is concave.

